# Molecular identification of a peroxidase gene controlling body size in the entomopathogenic nematode *Steinernema hermaphroditum*

**DOI:** 10.1101/2023.11.07.566113

**Authors:** Hillel T. Schwartz, Chieh-Hsiang Tan, Jackeline Peraza, Krystal Louise T. Raymundo, Paul W. Sternberg

## Abstract

The entomopathogenic nematode *Steinernema hermaphroditum* was recently rediscovered and is being developed as a genetically tractable experimental system for the study of previously unexplored biology, including parasitism of its insect hosts and mutualism with its bacterial endosymbiont *Xenorhabdus griffiniae*. Through whole-genome re-sequencing and genetic mapping we have for the first time molecularly identified the gene responsible for a mutationally defined phenotypic locus in an entomopathogenic nematode. In the process we observed an unexpected mutational spectrum following EMS mutagenesis in this species. We find that the ortholog of the essential *C. elegans* peroxidase gene *skpo-2* controls body size and shape in *S. hermaphroditum*. We confirmed this identification by inactivating the gene using CRISPR-Cas9. We propose that the identification of *skpo-2* will accelerate gene targeting in other *Steinernema* entomopathogenic nematodes used commercially in pest control, as *skpo-2* is X-linked and males hemizygous for loss of its function can mate, making *skpo-2* an easily recognized and maintained marker for use in co-CRISPR.

## Introduction

Entomopathogenic nematodes of the genera *Steinernema* and *Heterorhabditis* reside in the soil as developmentally arrested dispersal-stage infective juvenile (IJ) larvae (reviewed in Dillman and Sternberg 2012; Schwartz 2015). Upon encountering a suitable insect host, an entomopathogenic nematode invades its body, molts, and resumes development; in the process the nematode releases endosymbiotic pathogenic bacteria from its intestine into its insect host. Together the nematode and its bacterial symbiote rapidly kill the insect and convert the carcass into an incubator for the nematode-bacterial pair. When the carcass has eventually been exhausted, a subsequent generation of IJs, each carrying their cargo of pathogenic bacteria, will eventually disperse to begin the process anew. The entomopathogenic nematode lifecycle offers an opportunity to study the development and behavior of parasitic nematodes and their interactions with their bacterial symbiotes and their insect prey, along with other aspects of their biology shared with, and aspects of their biology differing from, those described in other nematodes.

A model for establishing entomopathogenic nematodes as a tool for laboratory research can be found in the extensive description of the biology of the free-living soil nematode *Caenorhabditis elegans*. Work on *C. elegans* has provided major contributions to our understanding of development and disease (Horvitz 2003; Sulston 2003; Brenner 2003; Fire 2007; Mello 2007). These studies have been particularly fruitful because of the features that make *C. elegans* a remarkably tractable organism in the laboratory: *C. elegans* is a small animal with a rapid generation time and reproduces by selfing hermaphroditism, all of which act to accelerate research using this organism, especially its molecular genetics (Apfeld and Alper 2018; Singh 2021). More recently, *C. elegans* was among the first animals to have their genome modified using CRISPR-Cas9 genome editing, opening new possibilities for the exploration of gene function (Frøkjær-Jensen 2013).

We are developing the entomopathogenic nematode *Steinernema hermaphroditum*, seeking to produce a similarly tractable and similarly powerful platform for laboratory research. This would enable research into aspects of its biology that are specific to this organism or that are found throughout entomopathogenic nematodes, but that are not amenable to study in any previously available nematode species, such as interactions between the nematodes and their bacterial symbiotes, interactions that are the product of millions of years of shared evolution. We have focused our efforts on the species *Steinernema hermaphroditum*. First reported in 2000 from field studies in the Moluccan islands of Indonesia and shown to be capable of reproducing for a single generation without mating, the species was subsequently lost for nearly twenty years before a 2019 report revealed it had been rediscovered outside New Delhi in 2019 (Griffin *et al*. 2000; Stock *et al*. 2004; Bhat *et al*. 2019). We have recently reported finding that this species consistently reproduces as a selfing hermaphrodite, established an inbred wild-type strain and protocols for its propagation in the laboratory, and showed it could be used in chemical mutagenesis screens to recover mutants that could then be used in complementation tests and genetically mapped against each other, and could be cryopreserved (Cao *et al*. 2022).

Continuing our efforts to develop this species as a platform for laboratory exploration, we sought proof-of-principle for molecular identification of a mutationally defined locus, which had not previously been achieved in an entomopathogenic nematode. Through whole-genome sequencing of three independent alleles of an X-linked locus causing a strong body size (Dumpy, or Dpy) phenotype we identified *Shm-skpo-2*, the *S. hermaphroditum* ortholog of the *C. elegans* peroxidase gene *Cel-skpo-2*, as the only mutated gene likely to be responsible for this Dpy phenotype. A mutation in *Shm-skpo-2* gene could not be separated recombinationally from the mutant phenotype, and the phenotype could be reiterated by knocking the gene out using CRISPR-Cas9 gene targeting. Loss of the *skpo-2* gene of *C. elegans* results in embryonic lethality; the difference in the mutant phenotypes of these two closely related genes highlights the importance of studying development in multiple distantly related species to more fully explore the range of phenotypes that can result from knocking out orthologous genes, making the functions of these genes more accessible in one species than in another. Because *Shm-skpo-2* loss-of-function mutants have an easily detectable mutant phenotype that is hemizygous in males and does not interfere with their ability to mate, we expect that the *skpo-2* orthologs of other *Steinernema* nematodes, including species already in use and in active development to control insect pests in agriculture, should serve as a useful marker for co-CRISPR experiments, making it possible to efficiently modify their genomes.

## Experimental procedures

### Nematode genetics

*Steinernema hermaphroditum* strains were derived from the inbred wild-type strain PS9179 and were cultured using the bacterial strains *Xenorhabdus griffiniae* HGB2511 and *Comamonas sp*. DA1877 as food sources essentially as described (Cao *et al*. 2022).

*Caenorhabditis elegans* were derived from the wild-type strain N2 and were cultured using *E. coli* OP50 as a food source as described (Brenner 1974). Existing *C. elegans* mutants obtained for use in this study included *skpo-1(ok1640) II* and *mlt-7(tm1794) II*, along with the balancer chromosome *tmC6[dpy-2(tmIs1189)] II*. Existing *S. hermaphroditum* mutants used included *dpy(sy1639) X*, *dpy(sy1644) X*, *dpy(sy1646) X*, *dpy(sy1662) X*, and *unc(sy1636) X*.

A genetic screen for visible phenotypic mutants of *S. hermaphroditum* was performed using ethyl methansulfonate (EMS) mutagenesis as described (Cao *et al*. 2022). A single phenotypic mutant, PS9839 *dpy(sy1926) X*, was recovered. Complementation tests were performed using *dpy(sy1926)* and the X-linked Dpy mutants *sy1646* and *sy1662*, marked with *unc(sy1636) X* to identify cross progeny.

### DNA sequencing and analysis

Genomic DNA was prepared essentially as described, except without grinding of frozen animals (Emmons *et al*. 1979). Animals were grown on 10 cm Petri plates containing NGM agar with a lawn of HGB2511 bacteria. Animals were washed repeatedly in M9 buffer and digested using proteinase K in the presence of SDS and beta-mercaptoethanol. Lysate was extracted with phenol/chloroform/isoamyl alcohol followed by chloroform. Nucleic acids were precipitated from the aqueous fraction using ethanol and recovered by spooling. RNA was removed by digestion with RNase A, after which DNA was recovered by ethanol precipitation. Purified DNA was sent to Novogene (Sacramento, CA) for Illumina sequencing with a target of 26.6 million paired-end 150 nt reads.

Analysis of high-throughput sequencing data was adapted from a published pipeline for *C. elegans* (Smith and Yun 2017). Sequencing reads were filtered using BBTools bbduk (http://sourceforge.net/projects/bbmap/) to remove reads matching an assembly of *X. griffiniae* HGB2511 genome sequence (Jennifer Heppert and Heidi Goodrich-Blair, personal communication). Reads were mapped to a draft annotated *S. hermaphroditum* PS9179 genome (Erich Schwarz, personal communication), reads were sorted, duplicate reads were removed, and reads were indexed using Samtools (Danecek *et al*. 2021). Mutations were detected using Freebayes (Garrison and Marth 2012) and were mapped onto gene models and categorized for coding changes using ANNOVAR (Wang *et al*. 2010). Annotated changes were sorted, compared, and counted using Excel (Microsoft, Redmond, WA).

Individual animals or small groups of animals were lysed and sequences were amplified from them using PCR essentially as described for *C. elegans* (Wicks *et al*. 2001) using oligonucleotide primers whose sequences are listed in Table S1. Restriction enzymes were obtained from New England Biolabs (Beverly, MA). For Sanger sequencing, at least two PCR products were combined for each sample; nucleic acid was purified using QiaQuick (QIAGEN, Germantown, Maryland) and sent to Laragen for Sanger sequencing (Laragen, Culver City, CA).

Homology searches of additional *Steinernema* nematodes were performed using BLAST 2.2.24+ on a Debian GNU server (Altschul *et al*. 1990) using genome and transcriptome assemblies downloaded from the NCBI or from WormBase ParaSite (Howe *et al*. 2017); accession numbers were *Steinernema carpocapsae* GCA_000757645.3 (DNA), *Steinernema carpocapsae* WBPS16 (mRNA), *Steinernema diaprepesi* GCA_013436035.1, *Steinernema feltiae* GCA_000757705.1, *Steinernema glaseri* GCA_000757755.1, *Steinernema hermaphroditum* GCA_030435675.1 (DNA and mRNA), *Steinernema khuongi* GCA_016648015.1, *Steinernema monticolum* GCA_000505645.1, and *Steinernema scapterisci* GCA_000757745.1 (Dillman *et al*. 2015; Serra *et al*. 2019; Baniya *et al*. 2019; Baniya and DiGennaro 2021). MEGA11 software (Tamura *et al*. 2021) was used to generate a neighbor-joining phylogeny using peroxidase protein sequences from *C. elegans* version WS290 (Davis *et al*. 2022) and from *S. hermaphroditum* GCA_030435675.1.

### CRISPR-Cas9

CRISPR-Cas9 targeting *skpo-2* in *C. elegans* was performed as described (Arribere *et al*. 2014) using *dpy-10* and *unc-58* co-conversion markers to obtain the two alleles *sy2121* and *sy2122*, respectively. In both cases the co-conversion marker was lost and the mutation was balanced using *tmC6[dpy-2(tmIs1189)]*. The guide RNA used contained the *C. elegans* genomic sequence CCCCAACATCGACCCATCTG. An oligonucleotide with the sequence CATCGGCGCCTACCCAGGCTATGACCCCAACATCGACCCATgggaagtttgtccagagcagaggtgactaagtgataagctagcCTGTGGCCAACGAGTTCACATCGTGCGCGTTCCGTTTTGG was included as a template for homology-directed repair of the double-strand break in *skpo-2* repair, including a STOP-IN cassette, in lowercase (Wang *et al*. 2018). Homology-directed repair was confirmed by Sanger sequencing. The *C. elegans* CRISPR protocol including its injection mixture was adapted for use in *Steinernema* with the exceptions that there was no co-conversion marker used and the injection mix was 1/10 Lipofectamine (Invitrogen, Waltham, MA). The guide RNA used included the *S. hermaphroditum* sequence GCACCCGAGGAAGGTACTCG and a repair template oligonucleotide with the sequence CATCGGCGCCTACCCAGGCTATGACCCCAACATCGACCCATgggaagtttgtccagagcagaggtgactaagtgataagctagcCTGTGGCCAACGAGTTCACATCGTGCGCGTTCCGTTTTGG, containing a STOP-IN cassette shown in lowercase, was included in the injection mix. Animals used for CRISPR-Cas9 genome editing were grown on DA1877 in preference to HGB2511.

Phenotypically Dpy F_1_ and F_2_ progeny of P_0_ animals injected with CRISPR reagents were recovered and used to establish clonal lines that were genotyped by PCR using the oligonucleotides GACGTGTGTTTCCTCCCGT and GCATCTTAGCCGGGAGACT followed by restriction digest with RsaI to detect changes at the CRISPR cleavage site and with NheI seeking evidence that the oligonucleotide template had been used as a template for homology-directed repair. One Dpy F_1_ animal contained two different alleles, with different PCR results at the *Shm-skpo-2* CRISPR target, that were segregated in subsequent generations.

Gas chromatography-mass spectrometry of an EMS sample was performed by Mona Shagholi of the Caltech Mass Spectrometry service center to confirm its molecular identity.

Strains are available upon request. Sequence data have been deposited in the NCBI Short Read Archive as part of NCBI BioProject PRJNA1037740.

## Results

### Molecular identification of a *Steinernema hermaphroditum dpy* gene by screens and sequencing

The first chemical mutagenesis screens in *S. hermaphroditum* used ethyl methanesulfonate (EMS) to recover 32 independent mutant strains with visible phenotypes such as uncoordinated (Unc) or dumpy (Dpy); mutants were identified as independent either because they had distinguishable phenotypes or because they arose from different mutagenized P_0_ animals (Cao *et al*. 2022). Three of these 32 mutations – PS9260 *dpy(sy1639),* PS9265 *dpy(sy1644)*, and PS9267 *dpy(sy1646)* – caused an identical Dumpy (Dpy) phenotype (Figure 1A) and were X-linked. These three mutations failed to complement each other, and genetic mapping showed each of the three was approximately 9 map units from the X-linked marker *unc(sy1635)*. Another X-linked mutation with a similar phenotype, *dpy(sy1662)*, complemented these mutations, indicating that *sy1639, sy1644*, and *sy1646* are in one complementation group and *sy1662* is in another.

**Figure 1.**
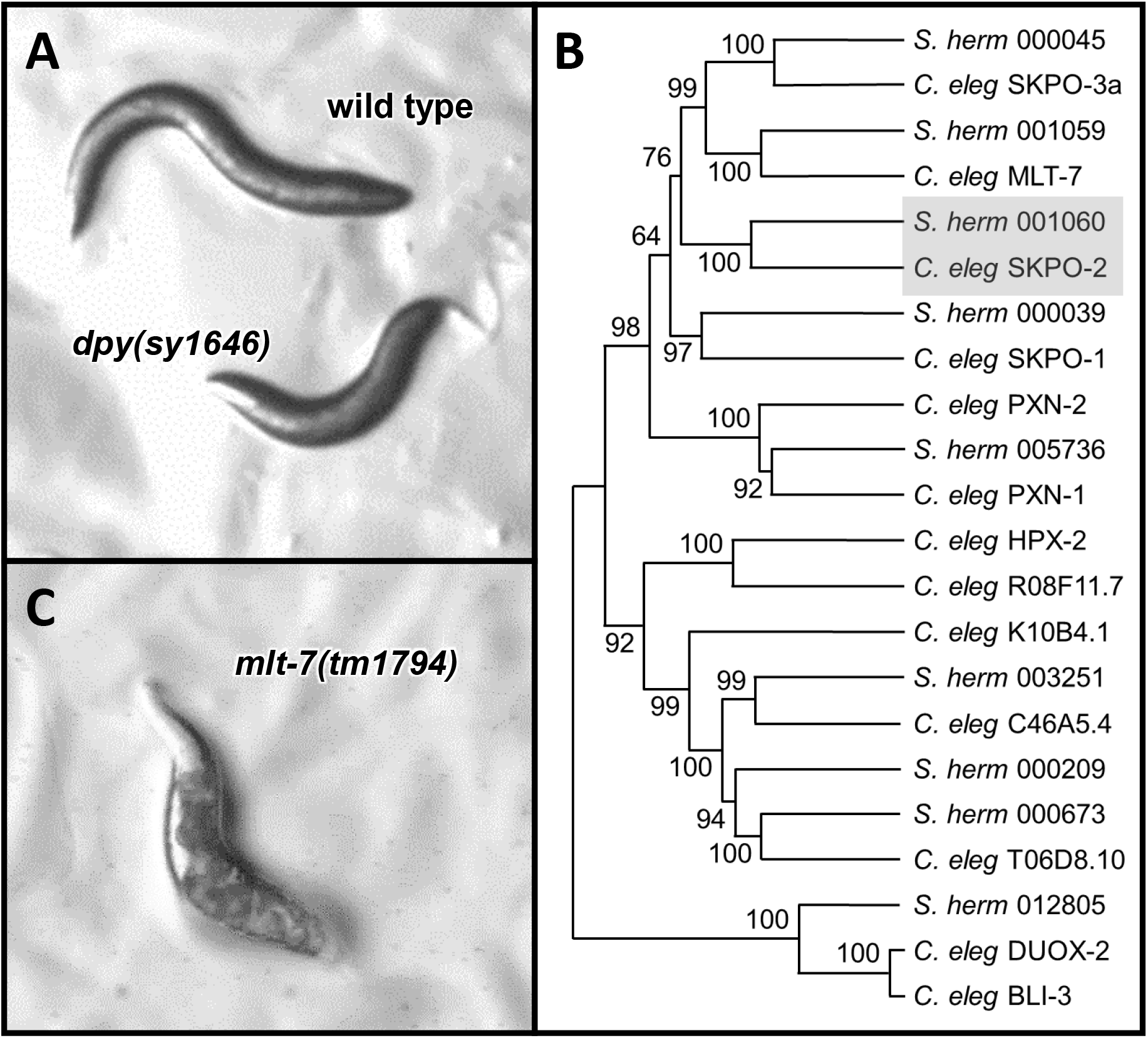
The Dpy phenotype of *S. hermaphroditum* and *C. elegans* peroxidase mutants. **A** Fourth-stage J4 larvae of the wild-type strain of *S. hermaphroditum* PS9179 and of the *skpo-2* mutant *sy1646*. **B** A phylogeny of *C. elegans* SKPO-2 and its closely related peroxidase proteins in *C. elegans* and *S. hermaphroditum*. *C. elegans* SKPO-2 and its *S. hermaphroditum* ortholog are indicated with a gray box. Branch strength bootstrap scores were generated using neighbor-joining with 1000 repetitions. *S. hermaphroditum* protein accession numbers are listed in Table S2. **C** The Dpy phenotype of rare surviving *C. elegans mlt-7(tm1794)* homozygotes. Although these rare survivors are generally healthy when their mother carried a wild-type allele, in subsequent generations the rare survivors are increasingly sickly and display pleiotropic phenotypes.

An additional round of EMS mutagenesis screens recovered one additional mutant, PS9839 *dpy(sy1926)*, with an indistinguishable Dpy phenotype. *dpy(sy1926)* was also X-linked and failed to complement *dpy(sy1646)* but did complement *dpy(sy1662)*, indicating *sy1926* was a fourth member of the complementation group containing *sy1639*, *sy1644*, and *sy1646*. We proceeded to whole-genome sequencing of three of these four allelic mutations: PS9260 *dpy(sy1639)*, PS9267 *dpy(sy1646)*, and PS9839 *dpy(sy1926)*. After reads had been filtered for bacterial contamination and mapped to a draft *S. hermaphroditum* genome and duplicate reads had been removed, we had 18.8x, 34.3x, and 46.0x coverage of the genomes of these mutants, respectively. We sought candidate genes that would be on the X-chromosome and would have coding mutations in all three strains; ideally these would be independent mutations, especially in the case of *sy1926*, recovered in a different screen from the first three alleles, and they were expected to be single-nucleotide C-to-T changes consistent with EMS mutagenesis (Anderson 1995; Volkova *et al*. 2020). These criteria resulted in four candidates: hypothetical proteins QR680_001060, QR680_001389, QR680_001390, and QR680_002483.

Further inspection suggested the latter three candidates were likely the result of sequencing and software issues: the mutations associated with these three candidates were defined by low read counts that had low quality scores. Proteins 001389 and 001390 are encoded by neighboring genes and include nearly identical sequence; these two genes have 14 different mutations annotated between them among the three strains, which did not seem consistent with the mutations having arisen after mutagenesis and being causative for the Dpy phenotype. Protein 002483 has 14 mutations annotated, of which three were annotated in more than one strain; this also is not consistent with the gene having been mutated to cause the Dpy phenotype. By contrast, the gene encoding protein 001060 has only three mutations annotated among the three strains, one in each strain; all three annotations have high read counts and quality scores.

We also examined the four multiply mutated X-linked genes’ homology to assess them as candidates. Predicted proteins 001389 and 001890 lack identifiable homologs by BLAST searches, with none found even in the other available *Steinernema* genomes or the *Steinernema carpocapsae* transcriptome, and lack conserved domains identifiable by SMART or by Pfam (Letunic *et al*. 2021; Mistry *et al*. 2021). The closest characterized homolog of protein 002483 is the product of the *C. elegans* gene *Cel-hgrs-1*; protein 002483 is also the predicted *S. hermaphroditum* protein most closely related to Cel-HGRS-1. RNAi-mediated inactivation of *Cel-hgrs-1* is reported to cause a Dpy phenotype among many other reported defects (Kamath *et al*. 2003), meaning this gene must be considered a viable candidate to cause the Dpy phenotypes of our *S. hermaphroditum* mutants, despite the low read count and the poor quality scores of the sequence data implicating this gene. The last of the four candidates is the gene encoding protein 001060, which is orthologous to the *C. elegans* gene *Cel-skpo-2*, encoding a predicted peroxidase (see Figure 1B). The mutant phenotype of *Cel-skpo-2* has not been reported, but it is closely related to *Cel-mlt-7*, loss of which causes defects in cuticle formation and molting along with nearly fully penetrant lethality and a Dpy phenotype in the rare animals that survive (Figure 1 C). Protein 001060 is more distantly related to the product of *Cel-bli-3*, which mutates to cause a blistered cuticle defect; *bli-3* is reported to function with *mlt-7* to regulate cuticle structure, and other blister mutants genetically interact with cuticular Dumpy phenotypes (Higgins and Hirsh 1977; Cox *et al*. 1980; Simmer *et al*. 2003; Thein *et al*. 2009). We have named the gene encoding protein 001060 *Shm-skpo-2* on the basis of its orthology to *Cel-skpo-2*.

Sanger sequencing confirmed the three mutations in *Shm-skpo-2* identified by high-throughput sequencing: PS9260 has a three-nucleotide deletion removing amino acid R503, PS9267 has a single-nucleotide G-to-T change causing the predicted coding change E469ochre; and PS9839 has a single-nucleotide G-to-T change causing the predicted coding change C178F (see Figure 2A). Attempts to identify a coding change in the fourth allelic *S. hermaphroditum* Dpy mutant, PS9265 *dpy(sy1644)*, which was not selected for whole-genome sequencing, demonstrated that the ninth exon of *Shm-skpo-2* could not be amplified using PCR primers that reliably amplified this sequence from the wild type, indicating that this strain contains a large deletion, insertion, or other rearrangement in this region of *Shm-skpo-2*.

**Figure 2.**
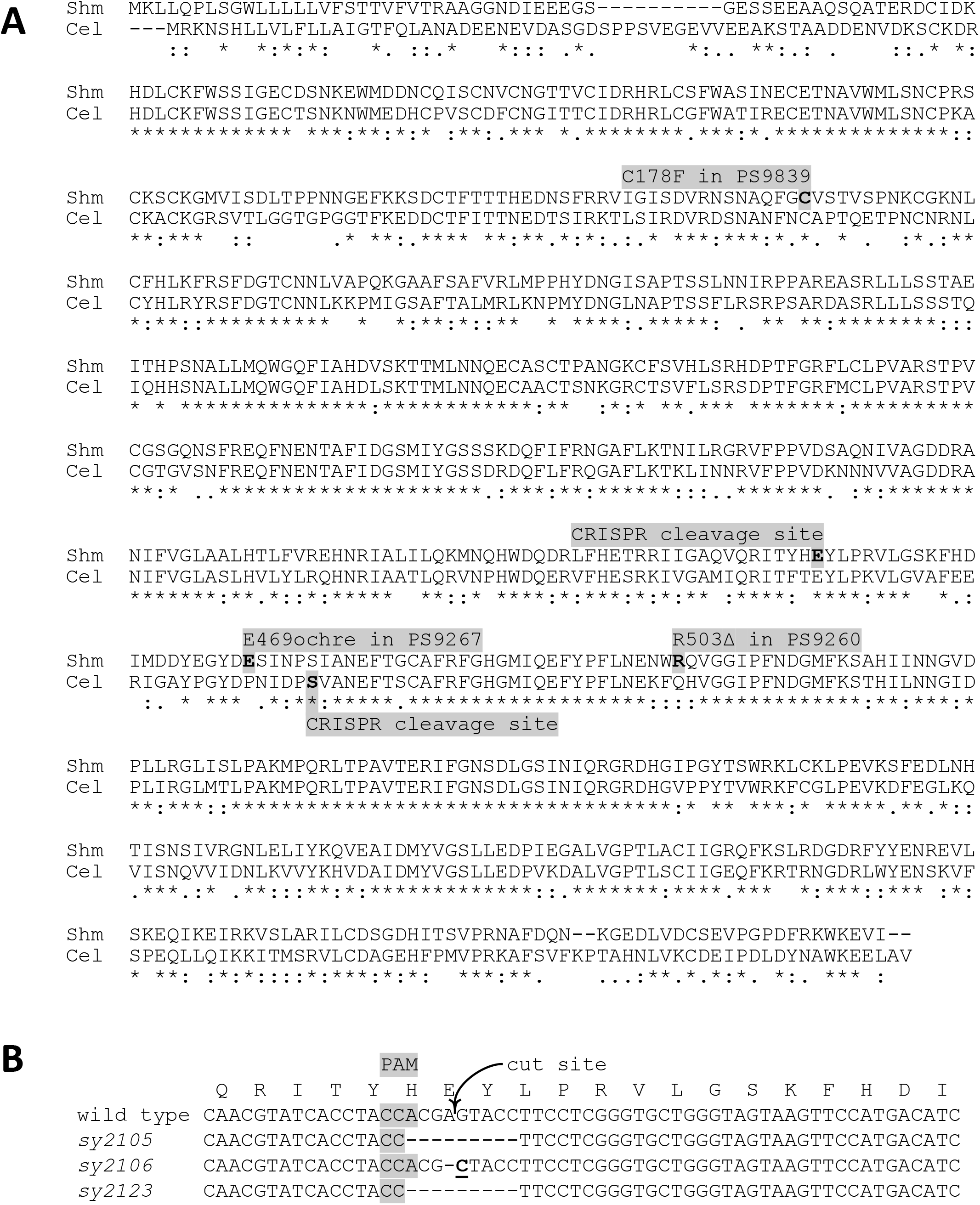
Mutations in *S. hermaphroditum* cause a Dpy phenotype. **A** An alignment of the orthologous SKPO-2 proteins of *S. hermaphroditum* (Shm) and *C. elegans* (Cel). The sites and effects of the three mutations detected by whole-genome sequencing of Dpy mutants are indicated, as are the sites targeted for cleavage by CRISPR-Cas9 in both *S. hermaphroditum* and *C. elegans*. Alignment and identification of conserved residues done by Clustal Omega as described in Materials and Methods; an asterisk (**\***) indicates an identical residue, a colon (:) indicates a highly similar residue, and a period (.) indicates a weakly similar residue. **B** identification of lesions in *S. hermaphroditum skpo-2* caused by CRISPR-Cas9. The three CRISPR-induced mutants whose targeted loci could be amplified by PCR were sequenced and found to cause small changes at the site targeted for cleavage.

One of the three molecularly identified mutations in *Shm-skpo-2*, the mutation in PS9839, disrupts a locally unique endogenous restriction site (FokI: GGATG). This restriction site was used to assess linkage between the *Shm-skpo-2* locus and the Dpy phenotype of PS9839: 0/118 Dpy self-progeny of *dpy(sy1926)*/+ heterozygotes contained wild-type sequence at *Shm-skpo-2*, indicating extremely tight linkage, within 2 map units at most (p<0.001). The *Cel-hgrs-1*-homologous gene encoding protein 002483 is nearly seven million base pairs from *Shm-skpo-2*, on a chromosome of approximately 18.4 million base pairs; tight linkage of the Dpy phenotype with *Shm-skpo-2* is inconsistent with the causative mutations being in the gene encoding protein 002483, leaving *Shm-skpo-2* as the only strong candidate identified by sequencing.

### Targeted inactivation of *Shm-skpo-2* using CRISPR-Cas9 causes a Dpy phenotype

To test the identification of *Shm-skpo-2* as the gene that mutates to cause the Dpy phenotypes of these four allelic mutations, we used CRISPR-Cas9 to knock out the gene’s function. A guide RNA was selected that should cause CRISPR-Cas9 to induce double-strand breaks within 65 nucleotides of the ochre stop mutation *sy1646* in PS9267. Six mutations were identified following CRISPR-Cas9 injection, each of which caused a stable, fully penetrant, healthy Dpy phenotype; all six were found to cause changes at the targeted site likely to disrupt gene function, either because they caused a genomic abnormality that prevented PCR of the *skpo-2* locus (*sy2107*, *sy2108*, and *sy2120*) or because they caused an alteration detectable by PCR and Sanger sequencing (*sy2015*, *sy2106*, and *sy2123*; see Figure 2B). Although an oligonucleotide donor was included as a template for homology-directed repair, the induced lesions were consistent with non-homologous end joining (NHEJ) (4/6 lesions) or with microhomology-mediated end joining (MMEJ) (*sy2015* and *sy2123*, which contain identical nine-nucleotide deletions that remove one copy of the directly repeated four-nucleotide sequence TACC and along with it the five intervening nucleotides before the next copy; *sy2105* and *sy2123* were isolated independently). Insertion of a STOP-IN cassette (Wang *et al*. 2018) at the corresponding site of the *C. elegans* ortholog *Cel-skpo-2* caused fully penetrant recessive embryonic lethality in that species; *trans-*heterozygotes between this lethal null mutation in *Cel-skpo-2* and the nearly lethal mutation in the closely related gene *Cel-mlt-7* were grossly wild type.

### EMS mutagenesis of *S. hermaphroditum* induced a mutational spectrum consistent with double-strand breaks

Of the four EMS-induced dumpy mutants in *Shm-skpo-2*, none were consistent with the expected mutational spectrum of EMS, which causes 95% single-nucleotide C-to-T substitutions in organisms ranging from the most extensively studied close relative of *S. hermaphroditum*, *C. elegans*, to the flowering plant *Arabidopsis thaliana* (Pastink *et al*. 1991; Greene *et al*. 2003; Volkova *et al*. 2020). A more comprehensive examination of the annotated sequence changes in each strain, focusing on those changes that were unique to each strain and should therefore have arisen subsequent to mutagenesis, showed that in each of the three mutant strains the single-nucleotide substitutions comprised roughly equal numbers of each nucleotide change (Figure 3A). The single-nucleotide substitutions accounted for approximately one-half to two-thirds of annotated mutations; approximately 15% of mutations were multi-nucleotide changes, and nearly all of the remaining 15-30% of annotated sequence changes were single-nucleotide insertions. The single-nucleotide insertions were almost exclusively found in noncoding sequences (not shown); as noncoding sequence is enriched for mononucleotide repeats, annotated single-nucleotide insertions might include spurious reports from the software used to detect mutations. The observed sequence changes were not consistent with mutagenesis using EMS nor with any other chemical mutagen causing single-nucleotide changes well characterized for mutational spectrum in nematodes, but were consistent with the mutations observed following the induction of double-strand breaks (Volkova *et al*. 2020). The mutations tended to appear in clusters rather than distributed evenly across the chromosome (Figure 3B). Mass spectrometry confirmed the chemical used to mutagenize was EMS, but as discussed below there are possible explanations for how EMS treatment could induce double-strand breaks instead of causing C-to-T single-nucleotide transitions.

**Figure 3.**
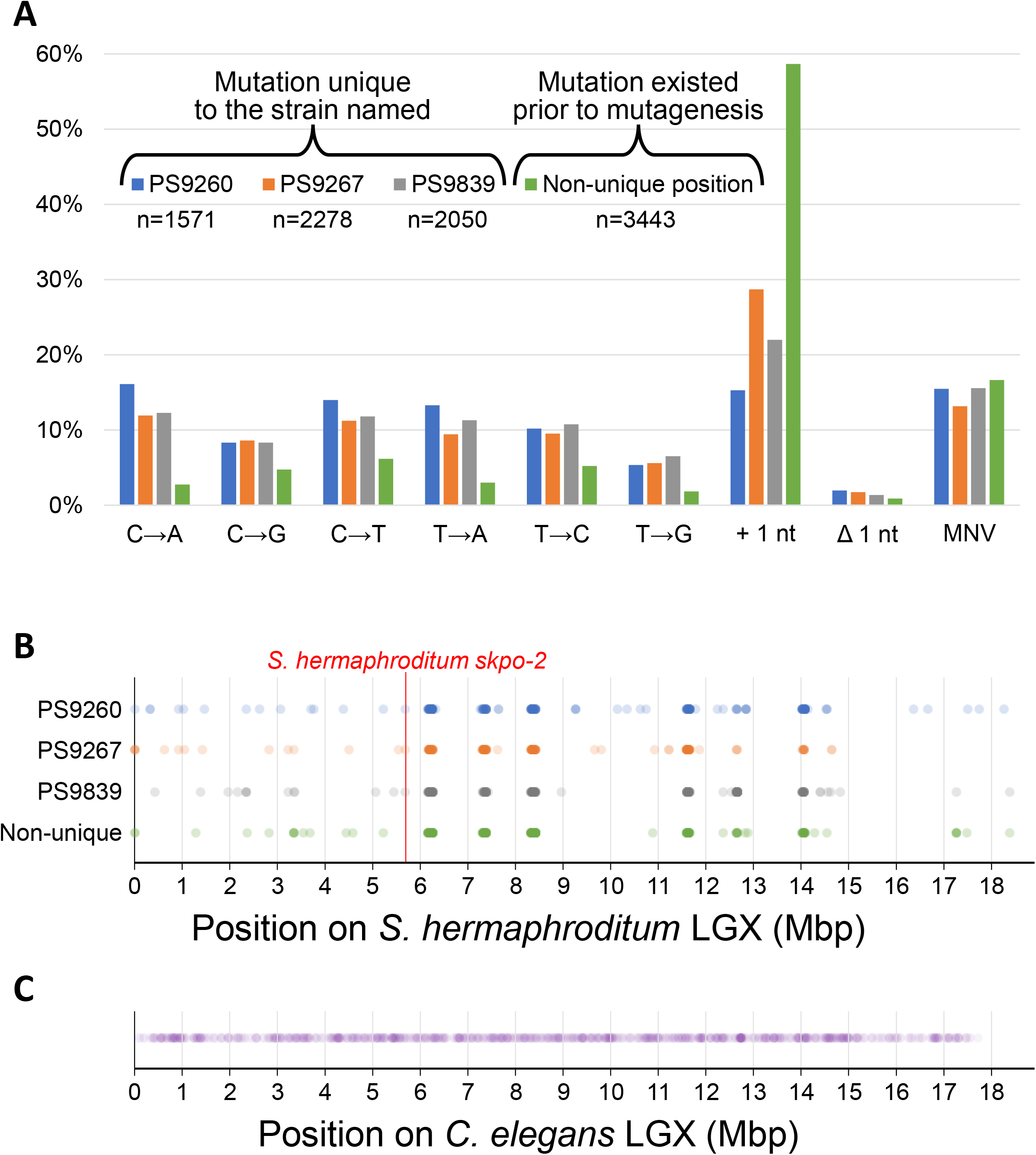
Mutational spectrum resulting from EMS mutagenesis of *S. hermaphroditum*. **A** The homozygous mutations found in the three sequenced strains were divided according to whether they were unique to the strain in question, or were found in multiple strains and so must have existed prior to mutagenesis. They were then sorted by the nature of the mutation: the single-nucleotide changes indicated, a single-nucleotide insertion, a single-nucleotide deletion, or a multi-nucleotide variation (MNV). The single-nucleotide insertion category was more common in pre-existing mutations, and almost all were intronic (not shown). The total numbers of unique mutations in each strain and the total number of shared mutations are indicated as “n=”. **B** Distribution of unique and common annotated mutations on the *S. hermaphroditum* X chromosome. Each row consists of from 227 to 295 mutations, each indicated with a colored, partially transparent circle; the intensely colored spots corresponding to hotspots for mutation detection indicate dozens of overlapping dots. The position of *skpo-2* on the chromosome is indicated. **C** Distribution of 868 EMS-induced mutations detected on the *C. elegans* X chromosome, from a collection of whole-genome sequencing data (HTS and PWS, manuscript in preparation). Each mutation is indicated with a colored, partially transparent circle. The distribution of EMS-induced mutations in *C. elegans* is noticeably more even than is seen in *S. hermaphroditum*.

## Discussion

### Molecular identification of a mutationally defined locus in an entomopathogenic nematode

The nematode worm *Caenorhabditis elegans* has been a uniquely powerful species for the use of experimental genetics to explore an animal’s biology (Horvitz and Sulston 1990; Meneely *et al*. 2019). *C. elegans* provides a model for other nematode species with similarly attractive characteristics suiting them to laboratory experimentation, whose biology differs interestingly from that of this free-living soil nematode. Entomopathogenic nematodes have several compelling aspects to their life cycle, offering experimental access to biology that has not been extensively explored in the laboratory. As one example, the relationships of entomopathogenic nematodes with their insect prey and with their pathogenic bacterial endosymbionts lack endogenous parallels in *C. elegans*; these relationships have important differences from those of *Pristionchus* nematodes, which offer a relatively well established experimental system but have a necromenic instead of pathogenic relationship with their insect hosts; they also differ from the commensal relationship between plant-pathogenic *Bursaphelenchus* nematodes, an emerging model system, and the insects that act as their vector (Sommer and McGaughran 2013; Félix *et al*. 2018; Kirino *et al*. 2023). We report our finding that the peroxidase gene *Shm-skpo-2* is required for normal body size and shape in *S. hermaphroditum*. This is the first time it has been possible to recover mutants of an entomopathogenic nematode and then to molecularly identify the gene responsible for preventing their phenotypic defect, and it is only the second time this has been reported for a clade IV nematode, following the recent molecular identification of a *tra-1* homolog mutated in sex determination mutants of *Bursaphelenchus okinawaensis* (Shinya *et al*. 2022).

*Shm-skpo-2* encodes a predicted peroxidase that is orthologous to the *C. elegans* gene *Cel-skpo-2* and is related to other homologous genes encoding predicted peroxidases. *skpo-2* and closely related genes are among several nematode genes closely related to the human peroxidasin PXDN (Thein *et al*. 2009). Human PXDN has been shown to crosslink collagen and regulate the structure of the endothelial basement membrane (Cheng *et al*. 2008; Bhave *et al*. 2012); peroxidase genes of *C. elegans* modify cuticle collagen structure and permeability, and many genes that mutate to cause or to suppress the Dpy phenotype in *C. elegans* encode collagens or proteins known to modify collagens (Edens *et al*. 2001; Myllyharju and Kivirikko 2004; Thein *et al*. 2009). It is likely that the Dpy phenotype of our *Steinernema* mutant is similarly the result of altered collagen structure, showing an evolutionarily conserved role of peroxidases in modifying and crosslinking collagens.

Loss of function of *Cel-mlt-7*, another *C. elegans* gene closely related to *Cel-skpo-2*, causes a Dpy phenotype not dissimilar from that found in mutants of *Shm-skpo-2* – but unlike *Shm-skpo-2* mutants, *Cel-mlt-7* mutants nearly all die during development, with only rare escapers surviving to show a Dpy phenotype, and the surviving Dpy animals are sick and show uncoordinated locomotion (Thein *et al*. 2009). *Cel-mlt-7* has an orthologous gene in *S. hermaphroditum*, distinct from *Shm-skpo-2*. The related but differing effects of mutating different homologs of peroxidasin in different nematodes is reflective of an established theme: although all nematodes share a highly similar body plan, the functions of orthologous genes can differ significantly even within a genus (Wang and Chamberlin 2002; Félix 2007; Mahalak *et al*. 2017), let alone among higher taxonomic units. Even if a particular phenotype has been extensively studied in one nematode, as the Dumpy phenotype has been in *C. elegans*, it can be expected that studies in a new, distantly related species will identify genes that cannot easily mutate to cause the phenotype in *C. elegans*, for example because of redundancy in that species or because pleiotropic phenotypes such as lethality prevent the recovery of such mutants, or because different genes are involved in the two species. Our studies in *Steinernema* have found viable, healthy Dpy mutants from a gene class that was known to be capable of modifying the cuticle, but that in *C. elegans* has thus far lacked for similarly healthy Dpy mutants; mutants of *Shm-skpo-2* could be studied to understand how these peroxidases affect cuticle structure and animal shape, in a way that the related *C. elegans* peroxidases cannot.

### An unexpected mutational spectrum following ethyl methanesulfonate mutagenesis

In the course of identifying *Shm-skpo-2* as the gene responsible for preventing the Dpy phenotype of four allelic mutants, we discovered that the mutations in *Shm-skpo-2* were not the single-nucleotide C-to-T changes expected from EMS mutagenesis, and indeed that the mutagenized strains each contained thousands of unique mutations, with a mutational spectrum that resembled those seen when double-strand breaks are induced rather than the spectrum normally expected following EMS mutagenesis; the mutational spectrum we observed also did not resemble the effects of mutagenesis with other chemicals causing single-nucleotide changes, nor did they resemble the observed effects of EMS mutagenesis on animals mutant for selected genes involved in DNA repair (Volkova *et al*. 2020). Our genetic screens for phenotypic mutants of *S. hermaphroditum* involved successful mutagenesis of the animals, and were not the result of spontaneous mutations: in the course of these screens we recovered dozens of stably phenotypic mutants in a very short time frame, from a small number of plates, and these mutants could be mapped and complementation tested – but phenotypic mutants were never seen spontaneously arising in the absence of a chemical mutagen, not even in the course of mapping, complementation testing, and cryopreserving our mutant collection, which involved propagating and inspecting vastly more animals than did the original screens. Treatment with the mutagen, which was confirmed to be EMS using mass spectrometry, must therefore have induced the genetic changes we detected, even though those changes did not conform to the mutational spectrum expected from EMS mutagenesis.

There was one other important clue: while EMS is noted for its ability to cause mutations that are distributed across the genome evenly (Figure 3C and Thompson *et al*. 2013), we recovered 33 phenotypic mutations of which four were alleles of one gene and two were alleles of another (Cao *et al*. 2022), suggesting that some loci were far more susceptible to our mutagenesis than others. Examination of the sites of the mutations we detected confirmed that the vast majority were tightly clustered at a few positions; these were the same positions in all three sequenced strains, and were also the sites of mutations found in common to the three strains, that must have predated mutagenesis. Between this clustering of mutations and the mutagenic spectrum we observed, we hypothesize that our mutagenesis treatment was inducing double-strand breaks, often at sensitive loci, rather than inducing single-nucleotide C-to-T transitions distributed evenly through the genome. An increased incidence of EMS induction of deletion mutations, presumed to be secondary to double-strand breaks, has previously been reported when an increased concentration of EMS was used to mutagenize *C. elegans* (Lesa 2006), suggesting that this result might reflect dose-responses for EMS mutagenesis differing between the two species; alternatively, EMS mutagenesis of cells in a state of cell-cycle arrest could cause double-strand breaks instead of C-to-T transitions. EMS mutagenesis normally converts cytosine residues to thymines when a guanine residue that has been modified by reaction with EMS to become O_6_-ethylguanine is misread during DNA replication and paired with thymine instead of cytosine (Sega 1984). If DNA replication were halted – if for example DNA checkpoint activity were different in *S. hermaphroditum* from what has previously been observed in *C. elegans*, or if the two species respond very differently to prolonged incubation in M9 buffer in the absence of food during the mutagenesis protocol – then the DNA base modifications caused by exposure to EMS would not rapidly be resolved to induce C-to-T single-nucleotide conversions, giving an opportunity for the altered residues to instead be repaired by error-prone nucleotide excision repair or for an accumulation of modified residues to trigger stalling of DNA replication followed by error-prone translesion repair of the clustered changes, or single-strand or double-strand breaks, whose resolution could result in a mutagenic spectrum characteristic of double-strand break repair (Kondo *et al*. 2010; Schärer 2013; Khatib *et al*. 2023). Further investigation of this difference between EMS mutagenesis of *S. hermaphroditum* and EMS mutagenesis of other nematodes using an essentially identical protocol should improve our ability to perform genetic screens probing the unique biology of this entomopathogenic nematode and may provide an opportunity to examine the basis of the differing effects of EMS mutagenesis on these different species.

### Prospects for CRISPR-Cas9 gene targeting in *Steinernema*

We confirmed our sequencing and linkage-based identification of *Shm-skpo-2* by using CRISPR-Cas9 to inactivate the gene, resulting in a collection of new mutants of this gene with an identical phenotype: healthy animals with a dumpy body shape. The resulting alleles were consistent with a mixture of NHEJ and MMEJ double-strand break repair (Xue and Greene 2021); we saw no evidence of homology-directed repair using the oligonucleotide template we had included. CRISPR-Cas9 is used extensively in established laboratory research animals and is emerging as a powerful tool in a growing variety of nematodes that have not previously been used extensively in laboratory research (Mendez *et al*. 2022; Cadd *et al*. 2022; Hellekes *et al*. 2023; Dutta *et al*. 2023); it is only with our work and with recent work from Mengyi Cao (personal communication) that CRISPR-Cas9 has been extended to entomopathogenic nematodes. We wish to particularly highlight the potential utility of the *skpo-2* locus in performing CRISPR studies using other *Steinernema* entomopathogens. Although we have focused our efforts on developing *S. hermaphroditum* as a tractable research system in the laboratory because its so-far unique reproductive biology of selfing hermaphroditism suits it unusually well to experimental genetics, every other *Steinernema* species whose reproduction has been described is diecious: they reproduce exclusively by outcrossing between males and females (Hunt and Nguyen 2016). Thus, any mutant phenotype used in these other *Steinernema* species must not interfere with the ability of phenotypic animals to mate, or else the mutant animals cannot easily be propagated. The availability of visible phenotypic markers is a matter of vital importance in using CRISPR-Cas9 for genome editing: it has been found in *C. elegans* that animals showing the phenotypic consequences of CRISPR-mediated genome modification at one locus are highly enriched for additional genome modifications at other sites simultaneously targeted using CRISPR (Kim *et al*. 2014). Variations of this method, called co-CRISPR and co-conversion, have been transformative for the efficiency of CRISPR-mediated genome modification in *C. elegans* and in other nematodes (Arribere *et al*. 2014; Cohen and Sternberg 2019; Choi and Villeneuve 2023). It is therefore desirable to have a *Steinernema* marker that can easily be recognized as identifying an animal that has had its genome modified using CRISPR, not only in *S. hermaphroditum* but also in other *Steinernema* species. The *skpo-2* locus is exceptionally well suited to serve as such as a marker: we could readily recover *Shm-skpo-2* mutants on the basis of their easily recognized phenotype, and *Shm-skpo-2* dumpy mutants are healthy and mate well as males. In each of the seven *Steinernema* species for which sequence is available, there is one *skpo-2* gene (Dillman *et al*. 2015; Serra *et al*. 2019; Baniya *et al*. 2019; Baniya and DiGennaro 2021). Because the *skpo-2* locus is X-linked in both *S. hermaphroditum* and *S. carpocapsae*, it is hemizygous in males; it is likely also X-linked in all other *Steinernema* nematodes, whose X chromosomes have not yet been identified. If loss of the *skpo-2* locus in other *Steinernema* nematodes has effects comparable to those we have observed in *S. hermaphroditum*, newly arising *skpo-2* mutant males in the first generation after CRISPR treatment should be dumpy, and yet be fully capable of mating, and will transmit any mutations they simultaneously carry in other loci targeted for genome modification using CRISPR. The *skpo-2* locus should therefore make it possible to efficiently modify the genomes of other *Steinernema* species already used commercially in agriculture to control insect pests (Karabörklü *et al*. 2017; Poinar 2018) and to probe the genomes of *Steinernema* species that possess novel biological abilities not yet observed or described in *S. hermaphroditum*, such as the abilities of *S. carpocapsae* to leap into the air and to secrete venom proteins into its insect host (Campbell and Kaya 1999; Lu *et al*. 2017; Dillman *et al*. 2021).

## Acknowledgments

We thank Erich Schwarz for generously providing early access to unpublished versions of the *S. hermaphroditum* genome and its annotation; Jennifer Heppert and Heidi Goodrich-Blair for unpublished HGB2511 sequence; Heenam Park and Tsui-Fen Chou for CRISPR reagents and advice about CRISPR; Mengyi Cao for unpublished information about *Steinernema* CRISPR; Barbara Perry, Wilber Palma, and Stephanie Nava for technical assistance; WormBase and WormBase ParaSite for *C. elegans* and *Steinernema* genome information; Mona Shagholi of the Caltech Mass Spectrometry service center; and Daniel Semlow and Anton Gartner for advice about the effects of EMS mutagenesis. Some strains were provided by the CGC, which is funded by the NIH Office of Research Infrastructure Programs (P40 OD010440). This work was supported by NSF-EDGE grant 2128267 (to PWS) and Caltech’s Center for Evolutionary Science (CES) and Center for Environmental Microbial Interactions (CEMI).

---

**Table S1.**
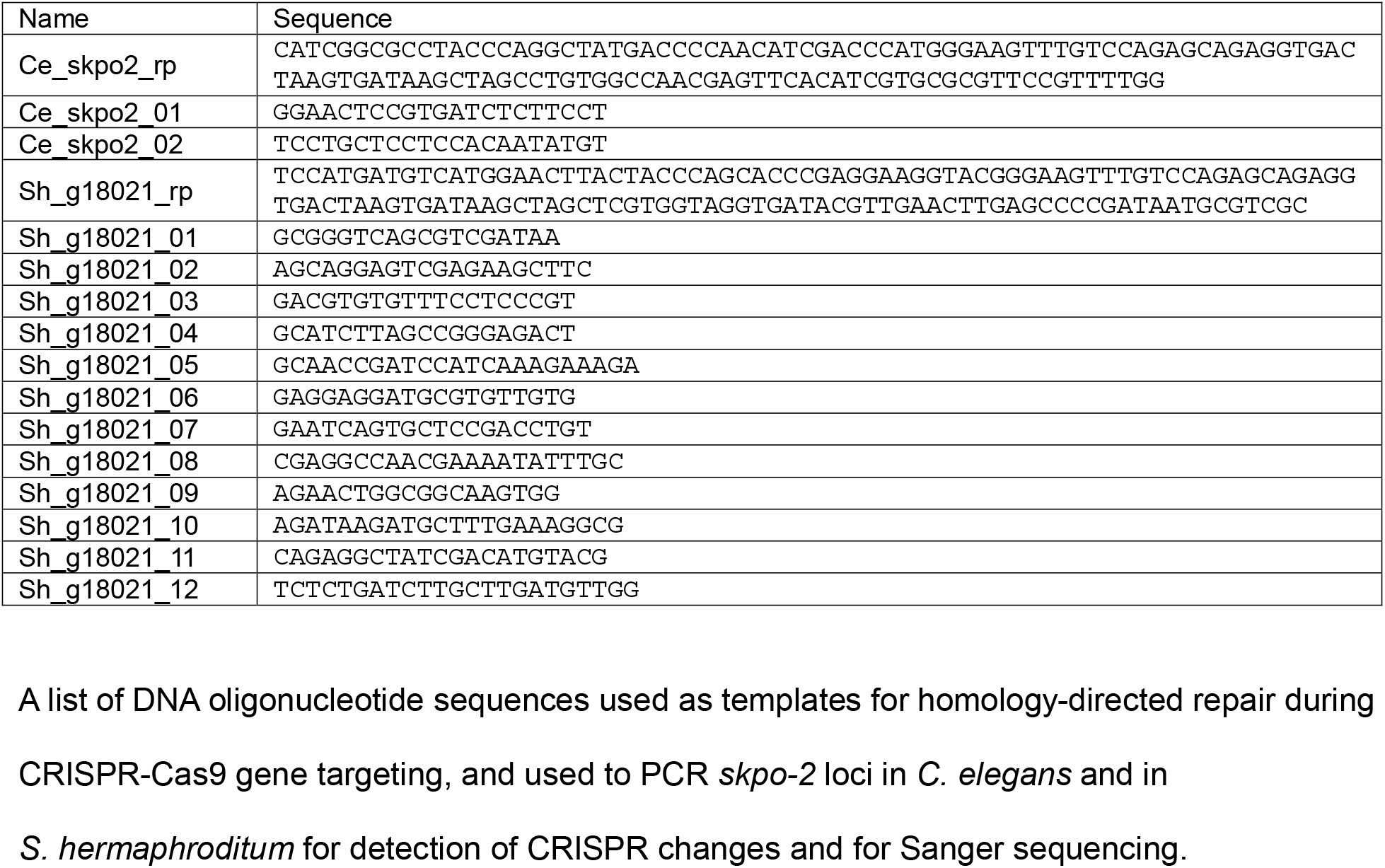
Oligonucleotides used in this study

**Table S2.** *Steinernema hermaphroditum* peroxidase protein accession numbers

| *S. hermaphroditum* protein | Closest *C. elegans* protein(s) | NCBI accession number |
| --- | --- | --- |
| QR680_000045 | SKPO-3a | KAK0393080.1 |
| QR680_001059 | MLT-7 | KAK0395007.1 |
| QR680_001060 | SKPO-2 | KAK0395009.1 |
| QR680_000039 | SKPO-1 | KAK0393066.1 |
| QR680_005736 | PXN-1 | KAK0411597.1 |
| QR680_003251 | C46A5.4 | KAK0399869.1 |
| QR680_000209 | T06D8.10 | KAK0393430.1 |
| QR680_000673 | T06D8.10 | KAK0394297.1 |
| QR680_012805 | DUOX-2, BLI-3 | KAK0417047.1 |

